# Matrix remodeling plays an etiological role in driving laminin-α2 deficient pathology

**DOI:** 10.64898/2026.06.28.735063

**Authors:** Veronica Pini, Anthony Accorsi, Ajay Kumar, Francesco Muntoni, Mahasweta Girgenrath

## Abstract

Laminin-α2 (gene: *LAMA2*) is a key protein in the basement membrane of muscle and Schwann cells. A complete lack of this protein results in LAMA2-related congenital muscular dystrophy (LAMA2-RD), a severe muscle disease characterized by progressive muscle weakness, respiratory insufficiency, failure to thrive and shortened life span. One key signature of this disease is early onset of fibrosis coupled with poor muscle growth. We previously showed that TGF-β and its activator, integrin-αV, are elevated in dystrophic fibers of Dy^W^ mice, a mouse model of LAMA2-RD. Other than activating TGF-β, integrin-αV is also known to facilitate the transdifferentiation of various cell types to myofibroblasts. In this study we present evidence for transcriptional dysregulation of genes driving myofibroblast transdifferentiation and extracellular matrix (ECM) remodelling during the early development of Dy^W^ mice that is also reflected in muscle biopsies from young LAMA2-RD patients. We hypothesize that the early ECM remodelling, seen in both Dy^W^ mice and LAMA2-RD children, may explain the congenital onset of fibrosis with poor muscle growth seen in the disease.

**Summary Statement:** Characterization of fibrogenic pathways that play potentially etiological roles in in LAMA2-related congenital muscular dystrophy

## Introduction

Congenital muscular dystrophies (CMDs) are a clinically and genetically heterogeneous group of diseases arising mostly from defects in proteins associated with the dystrophin-glycoprotein complex at the muscle cell membrane. Of these, mutations in the α2 chain of the heterotrimeric laminin-211 protein result in LAMA2-related CMD (also known as LAMA2-RD)^1–3^. This protein is predominantly found in the basement membrane of skeletal muscle and Schwann cell. In the extracellular matrix, it interacts with nidogen and agrin, while at the sarcolemma its cognate binding partners are α-dystroglycan and α7β1-integrin. The absence of this protein results in impaired anchoring of myofibers and signalling dysregulation, leading to severe downstream pathologies such as chronic inflammation, widespread fibrosis, progressive muscle wasting, and diminished regeneration^4–6^. The disease presents at or near birth, and children with LAMA2-RD typically exhibit severe hypotonia and muscle weakness, and almost never achieve independent ambulation. These patients often face significantly shortened life expectancy, primarily due to respiratory insufficiency and failure to thrive, which are two major drivers of morbidity and mortality ^1,7–9^. Despite extensive research into the disease’s underlying mechanisms, there is currently no cure or effective treatment for LAMA2-RD, highlighting a significant unmet need for this patient population.

While many disease drivers have been elucidated in the Dy^W^ mouse model of LAMA2-RD, early, excessive and diffused fibrosis, with non-structural ECM protein (matricellular proteins) dysregulation standing out as distinctive and possible driving characteristic of this disease^10^. Although TGF-β is known to play a major role in the induction and progression of fibrosis^11–17^, much remains to be clarified about the other interconnected signaling pathways that contribute to the severe fibrotic pathology observed in Dy^W^ mice and patients.

One area of convergence between matricellular protein dysregulation and TGF-β is integrin signaling, specifically through integrin-αV^17–24^. Integrins are heterodimeric membrane-spanning proteins that are involved in numerous critical cellular functions including adhesion and signal transduction^25^. Dysregulation of integrin expression and/or signaling has been implicated in numerous diseases including cancer^26^, heart disease^27^, and the fibrosis of various organs^19^. Specifically, integrin-αV is known to activate TGF-β by liberating it from its latent complex in the ECM thereby allowing it to bind to its cognate cell surface receptors and initiate downstream signaling^21^. We showed previously that integrin-αV, its beta dimer partners, and active TGF-β were all upregulated in adult Dy^W^ mice^28^. Further, we showed that integrin-αV was expressed at the sarcolemma of myofibers in adult Dy^W^ mice, a previously unappreciated pathological phenomenon. We also showed that treatment with the Angiotensin II type 1 receptor blocker Losartan not only alleviated fibrosis in these mice but also reduced the expression of integrin-αV and circulating activated Tgf-β^28^. Together, these findings point to the possibility that integrin-αV may be playing a critical role in driving fibrosis in Dy^W^ mice.

A further consequence of the reciprocal interplay between integrin-αV and TGF-β is the induction of myofibroblast transdifferentiation in several cell types^16,29–33^. TGF-β signaling stimulates the expression of epithelial to mesenchymal transdifferentiation (EMT)-related transcription factors (Slug, Snail, Twist), which then facilitate changes in gene expression and in the integrin isoform repertoire to facilitate myofibroblast transdifferentiation (MT)^16,17,30,33–35^. Myofibroblasts are characterized as hyperproliferative and producing more ECM material than quiescent tissue-resident fibroblasts^36–43^. Additionally, the myofibroblasts are hypercontractile (due to the induction of α-smooth muscle actin stress fibers) and can increase ECM stiffness which is not only inherently detrimental to normal muscle function but can also be a positive regulator of integrin-TGF-β interactions. This interplay of signaling pathways create a feed-forward loop driving matrix remodelling and myofibrogenesis^17,20,31,38–40,44^.

In this study we established for the first time the key role of TGF-β and integrin-αV during early postnatal development in Dy^W^ mouse. Importantly, we observed that the dysregulation of fibrotic pathway genes documented in the Dy^w^ mouse is recapitulated in the biopsies of affected patients with LAMA2-RD. The extensive changes observed in the myomatrix early in life could therefore explain the derailed myogenesis and poor muscle growth responsible for the severe phenotype observed in LAMA2-RD patients. Lastly, we showed here that antifibrotic intervention such as Losartan treatment reverses integrin expression and myofibrogenesis protein upregulation. This work further validates integrin and TGF-β signaling pathways as integral contributors to muscle fibrosis, the significant potential of therapies that will target fibrosis in LAMA2-RD, as well as elucidation of potential biomarkers for fibrotic resolution.

## Results

### Myofibroblast-related proteins are upregulated in adult Dy^W^ mice

The TGF-β-Integrin-αV signaling is intricately involved in the transdifferentiation of various cell types to a myofibroblast phenotype^16,19,21,30^. Based on the significant upregulation of matricellular proteins, integrin-αV and TGF-β signaling in adult Dy^W^ muscle^28^, which all feed into MT, we examined if there was any evidence of MT in adult Dy^W^ muscle. We examined the proliferative properties of fibroblasts isolated from hind leg muscles (quadriceps, tibialis anterior and gastrocnemius/soleus complex) of young adult (5-week old) Dy^W^ mice in comparison to fibroblasts isolated from WT muscle at the same time point (n=4 animals). We found that not only there are significantly more fibroblasts per milligram of muscle tissue in Dy^W^ muscle but, when plated at equal densities, Dy^W^ fibroblasts are significantly (p-value <0.05) more proliferative than WT fibroblasts, a behavior more like myofibroblasts (*Fig. 1*). Of note, this hyperproliferative nature subsides after a few passages in culture (personal observation), suggesting that the dystrophic environment of Dy^W^ muscle may prime these cells towards the myofibroblast phenotype.

**Fig. 1.**
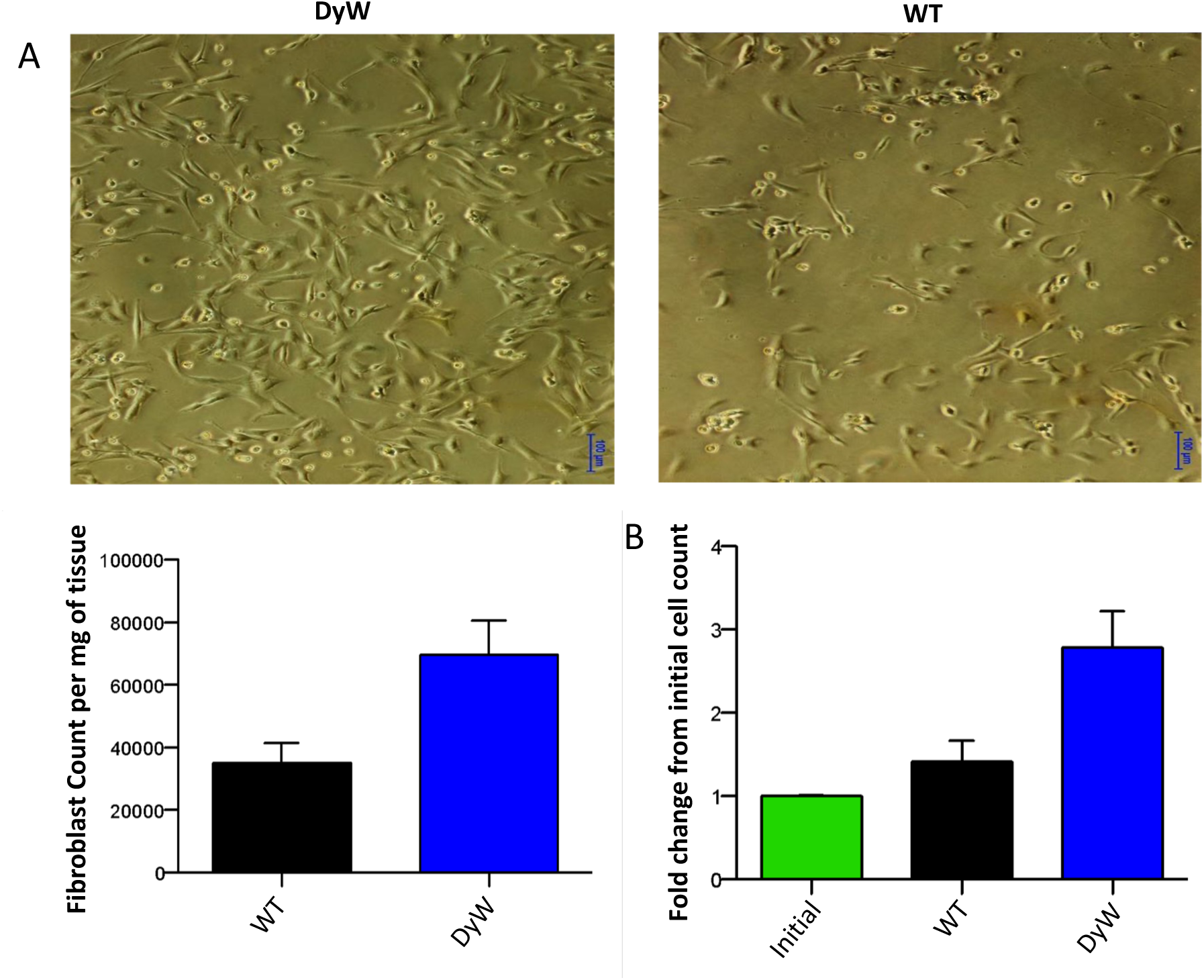
Dy^W^ fibroblasts are hyperproliferative in culture. A) Isolation of fibroblasts from Dy^W^ and WT muscle tissue show there are more fibroblasts per mg of muscle tissue in Dy^W^ muscle. B) When plated at equal density, Dy^W^ fibroblasts are also hyperproliferative compared to WT fibroblasts.

A classical feature of myofibroblasts is the induction of alpha-smooth muscle actin (α-SMA) stress fibers that imparts the ability to contract the surrounding matrix. Other proteins involved in the trans-differentiation and identification of myofibroblasts include the intermediate filament protein vimentin (Vim), the platelet derived growth factor receptor beta (Pdgfr-β), and osteoblast cadherin (Cdh11) which allows myofibroblasts to adhere to the stiff matrices they produce. We found that genes coding for these proteins were upregulated in dystrophic hindlimb muscles, as shown from their increased transcript and protein (p<0.01, n=4) as early as seven-weeks old Dy^W^ mice (Fig. 2A-D). We saw numerous mononuclear α-SMA positive cells in Dy^W^ muscle, some of which appeared to be mature myofibroblasts as evidenced by the expression of desmin^45^. We also observed an expansion of vimentin-positive cells in the interstitium of tibialis anterior muscles isolated from Dy^W^ mice (Fig. 2G).

**Fig. 2.**
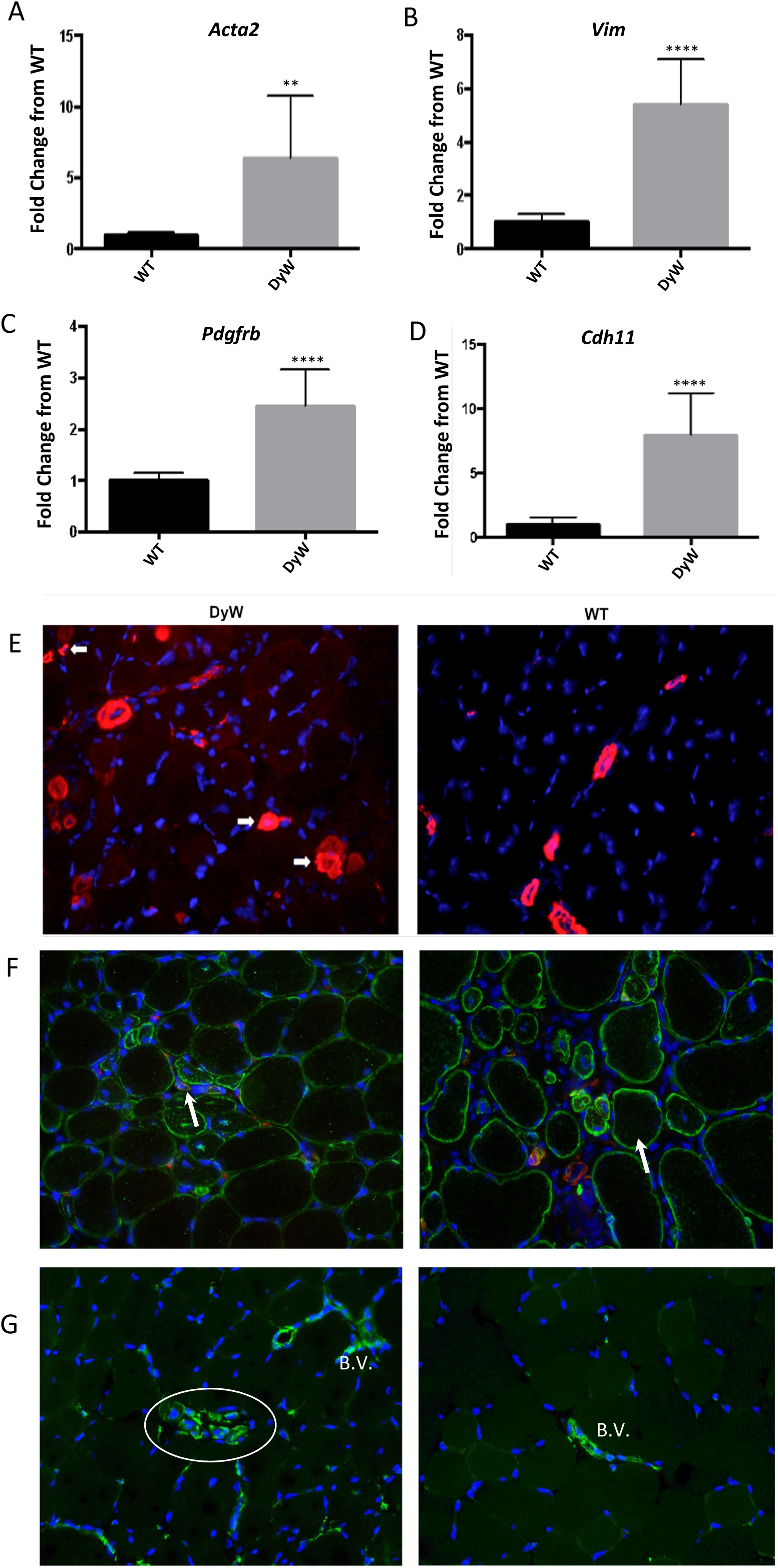
Myofibroblast markers are upregulated in adult Dy^W^ mice. A-D) Gene expression measured by qRT-PCR shows significant upregulation (p<0.01, n=4, t-test) of α-smooth muscle actin (A), vimentin, (B), platelet derived growth factor receptor beta (C), and osteoblast cadherin (D). **=p<0.01, ****=p<0.0001. E) Staining with anti-α-SMA antibody shows multiple mononuclear α-SMA positive cells (arrows) in Dy^W^ muscle tissue that are not apparent in WT muscle. α-SMA staining appears localized to blood vessels in WT mice. F) Co-staining with anti-α-SMA and anti-desmin antibodies shows that many of the α-SMA positive cells are also expressing the intermediate filament desmin, further supporting they are myofibroblasts. G) Immmunostaining with anti-vimentin antibody shows vimentin positive mononuclear cells not associated with blood vessels. WT tissue only shows vimentin expression around blood vessels. B.V=blood vessel.

We next assessed the gene expression levels of transcription factors downstream of Tgf-β that are responsible for MT in other disease scenarios, namely Snail, Slug, and Twist as well as β-catenin and one of its fibrosis-promoting cofactors (*Tcf4*). We found that at seven weeks of age, hindlimb muscles of Dy^W^ mice have significantly elevated transcript levels (p<0.05, n=4) of all these proteins compared to tissue obtained from age-matched WT mice (Fig. 3A-E). Protein evaluation via immunostaining of Snail, Slug and β−Catenin showed a corresponding increase in protein levels in Dy^W^ mice (Fig. 3F-H). Taken together, the hyperproliferation of Dy^W^ fibroblasts coupled with significant upregulation of the myofibroblast marker proteins and transcription factors seen in Dy^W^ muscle tissue provides considerable evidence for the presence of myofibroblast trans-differentiation in Dy^W^ mice that could be playing critical role in driving disease progression.

**Fig. 3.**
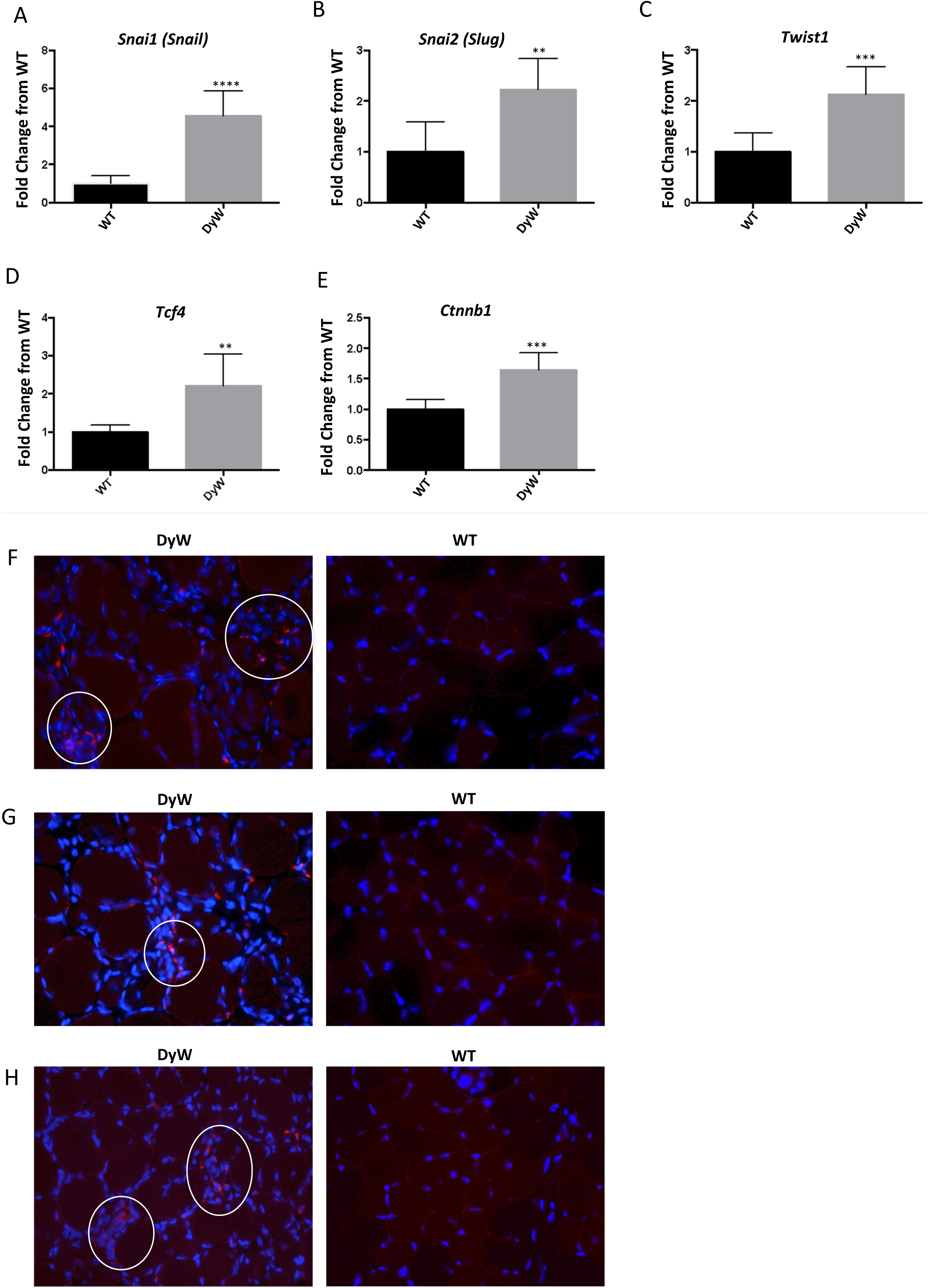
Myofibroblast-related transcription factors are upregulated in Dy^W^ muscle. A-E) Gene expression measured by qRT-PCR shows significant upregulation (p<0.01, n=4, t-test) of transcription factors known to govern myofibroblast transdifferentiation including the transcription factors Snail (A), Slug (B), and Twist1 (C), as well as Tcf4 (D) and b-catenin (E). F-H) Immunostaining with anti-snail (F), anti-slug (G), and anti-β-catenin (H) antibodies shows cells positive for these transcription factors in Dy^W^ muscle but not in WT muscle.

### Integrins, growth factors, and proteins related to myofibrogenesis are dysregulated throughout early development of Dy^W^ mice

As LAMA2-RD is a congenital disease and presents at/near birth, the most relevant data concerning disease evolution comes from early time points. Following the elucidation of integrin dysregulation and the presence of myofibroblast transdifferentiation in adult Dy^W^ mice, we next examined if these altered pathways could potentially be etiological to Dy^W^ pathology and thus similarly drive pathology in LAMA2-RD patients. Gene expression, via mRNA transcripts levels of α and β integrins as well as a relevant downstream effector protein integrin-linked kinase (*Ilk*) were measured from pooled hind limb muscle of Dy^W^ and WT mice collected at postnatal weeks one, two, three, and four (Fig. 4). Integrin-αV, previously mentioned to be a critical mediator of pathological Tgf-β activation, was upregulated (p<0.05, n=4) in Dy^W^ mice over age-matched WT mice at all time points. In addition, we found that integrin-α5, also known as the fibronectin receptor playing roles in macrophage recruitment and NF-κB signaling^46^, was also upregulated (p<0.05, n=4) throughout development, along with key integrin β dimer partners and adaptor kinases. Along with integrin expression, we also examined transcript expression of relevant growth factors known to be downstream of integrin activation and drive the fibrotic pathology (Fig. 5). We found significant dysregulation of the two Tgf-β isoforms known to be reliant on RGD-mediated binding of the latency-associated peptide (LAP) binding for activation (Tgf-β1/3)^22,24,47^.

**Fig. 4.**
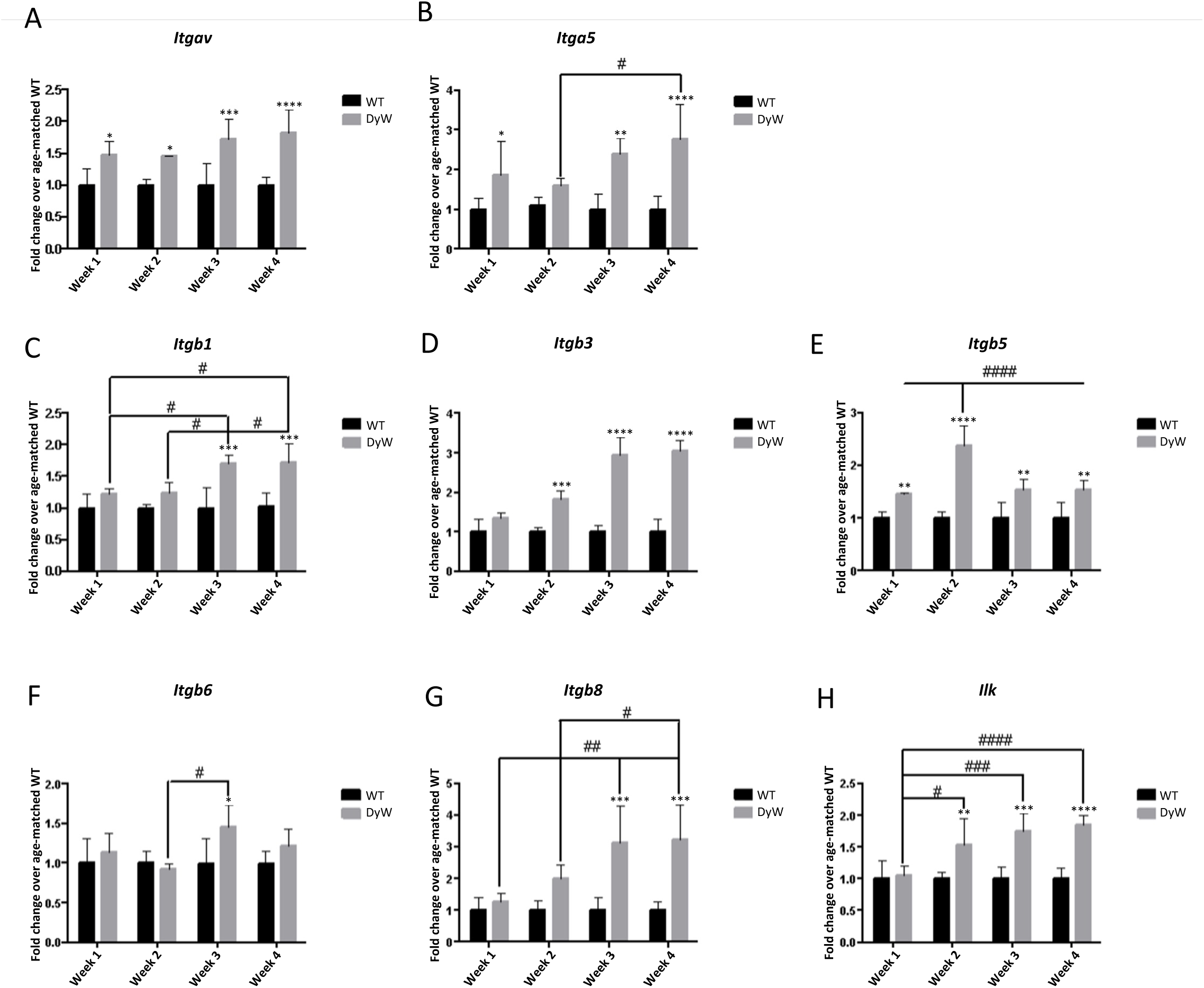
Integrin subunits are dysregulated during Dy^W^ development. A-H) Gene expression measured by qRT-PCR show that Integrin-αV, as well as Integrin-α5, and β subunits known to pair with Integrin-αV and activate Tgf-β are upregulated throughout Dy^W^ development. All subunits are significantly upregulated (p<0.05, n=4, two-way ANOVA) in weeks 3 and 4 with the exception of Integrin-β6 at week 4. Key signaling protein downstream of Integrin-αV, Integrin-linked kinase, is also significantly (p<0.001, n=4, two-way ANOVA) upregulated in weeks 2-4 * denotes significance between age-matched Dy^W^ and WT. # denotes significance between Dy^W^ groups. *=p<0.05, **=p<0.01, ***=p<0.001, ****=p<0.0001. Trend applies to # as well.

**Fig. 5.**
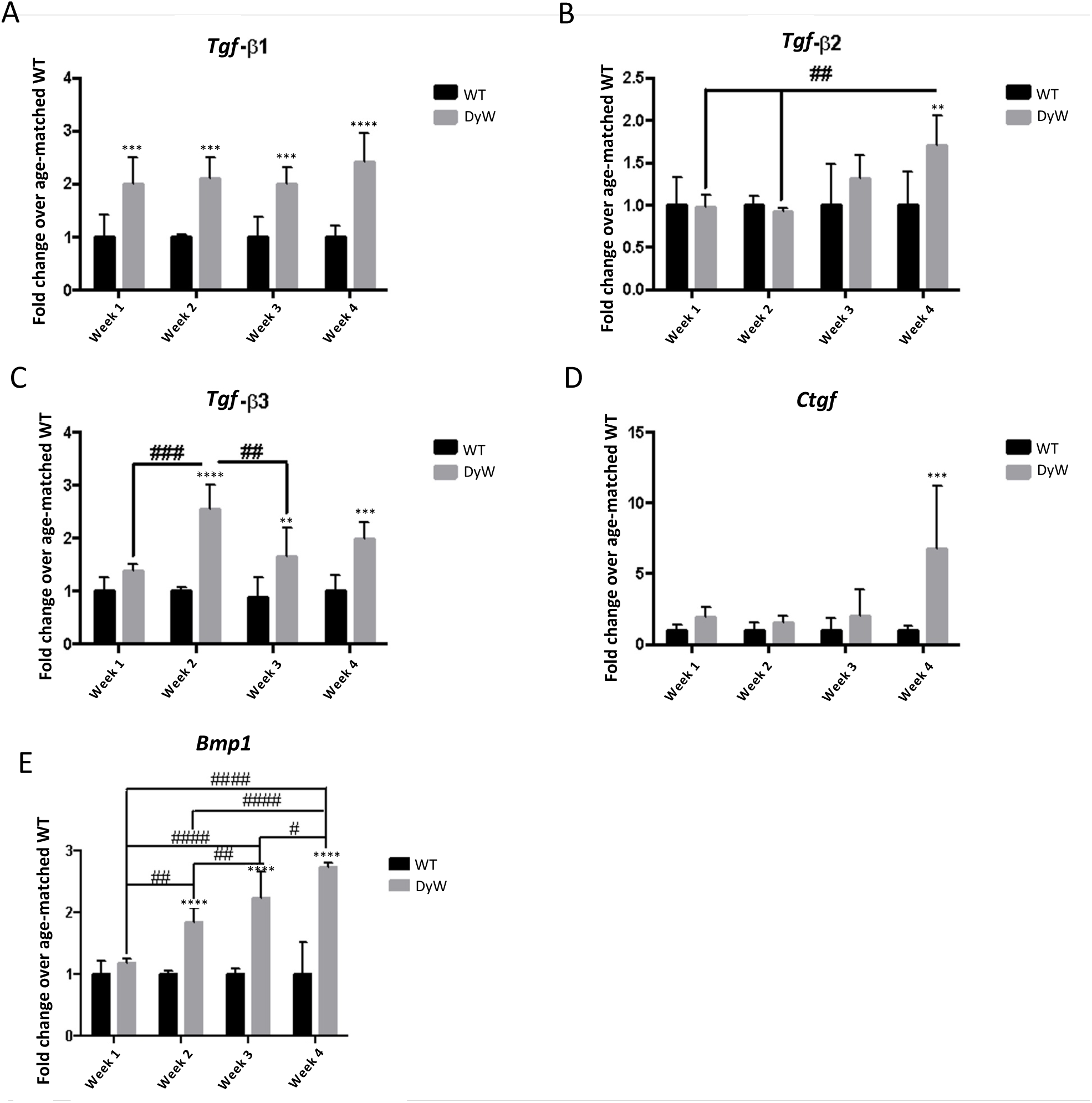
Fibrotic growth factors are dysregulated throughout Dy^W^ development. A-E) Gene expression measured by qRT-PCR shows that growth factors known to drive fibrosis including Tgf-β isoforms 1 (A), 2 (B), and 3 (C), Ctgf (D), and Bmp1 (E) are significantly dysregulated (p<0.01, n=4, two-way ANOVA) throughout Dy^W^ postnatal development. Interestingly, Tgf-β isoforms 1 and 3 show most dysregulation and are the two isoforms that have RGD sequences in their LAP and therefore are the only ones that can be activated by Integrin-αV. * denotes significance between age-matched Dy^W^ and WT. # denotes significance between Dy^W^ groups. *=p<0.05, **=p<0.01, ***=p<0.001, ****=p<0.0001. Trend applies to # as well.

We also examined transcript levels of numerous other growth factors (Fig. 5), matrix proteins (Fig. 6), cytoskeletal proteins (Fig. 6), and transcription factors (Fig. 7) known to be associated with this pathway. We found many of these genes to be upregulated in Dy^W^ muscle compared to age-matched WT mice throughout early development. Taken together, the aggregate results point to an early dysregulation of proteins known to drive myofibrogenesis and myomatrix remodeling. These may create a microenvironment suitable for further fibrotic development that is inhibitory toward efficient postnatal myogenesis, as seen in Dy^W^ mice^10^.

**Fig. 6.**
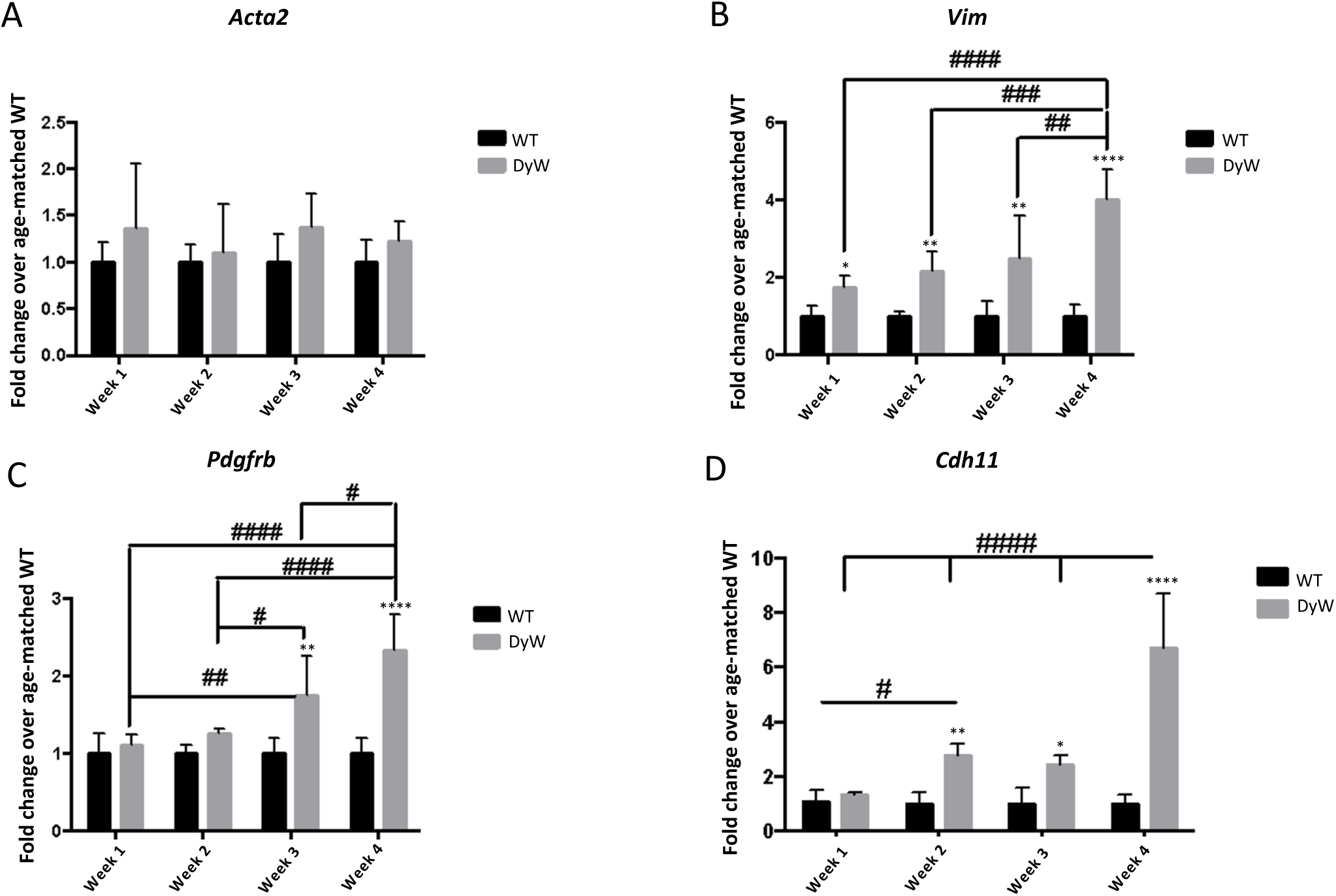
Myofibroblast markers are dysregulated in developing Dy^W^ mice. A-D) Gene expression measured by qRT-PCR shows significant dysregulation (p<0.05, n=4, two-way ANOVA) of proteins found on myofibroblasts including vimentin (B), platelet derived growth factor receptor beta (C), and osteoblast cadherin (D). Interestingly, α-SMA is not transcriptionally upregulated during Dy^W^ development (A). * denotes significance between age-matched Dy^W^ and WT. # denotes significance between Dy^W^ groups. *=p<0.05, **=p<0.01, ***=p<0.001, ****=p<0.0001. Trend applies to # as well.

**Fig. 7.**
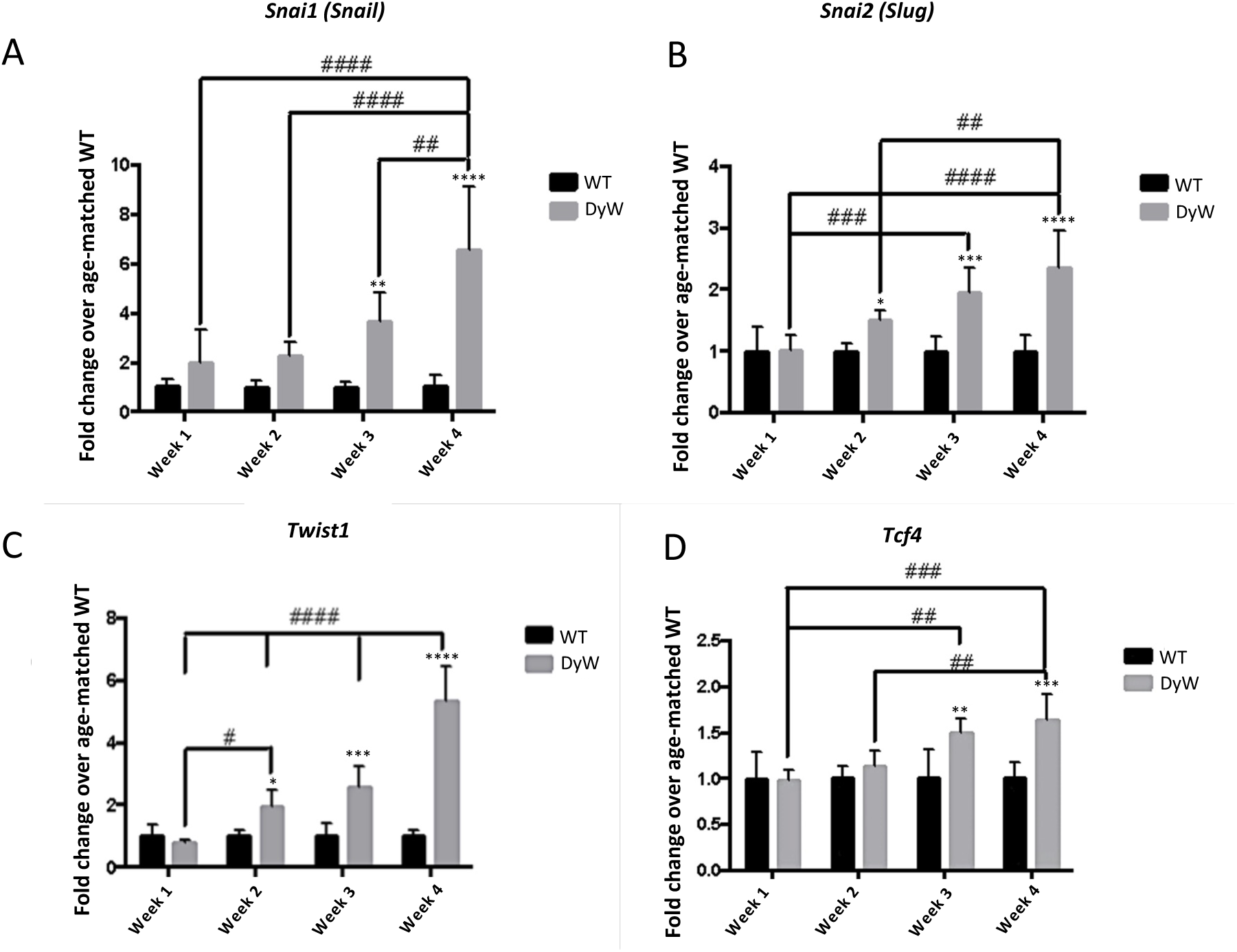
Myofibroblast transdifferentiation transcription factors are dysregulated throughout Dy^W^ development. A-D) Gene expression measured by qRT-PCR show significant dysregulation (p<0.01, n=4, two-way ANOVA) of transcription factors known to drive myofibroblast transdifferentiation including the bHLH transcription factors snail (A), slug (B), twist1 (C), as well as TCF4 (D) in postnatal weeks 2-4. * denotes significance between age-matched Dy^W^ and WT. # denotes significance between Dy^W^ groups. *=p<0.05, **=p<0.01, ***=p<0.001, ****=p<0.0001. Trend applies to # as well.

### Evidence of matricellular protein dysregulation and myofibrogenesis in LAMA2-related congenital muscular dystrophy patient biopsies

To investigate if murine findings are conserved in patients, we assessed an array of fibrosis-related genes in quadriceps biopsies obtained from patients with genetically confirmed LAMA2-RD against age-matched controls that presented with hypotonia but in whom we had eventually excluded a neuromuscular disease. We found that many of the genes seen to be upregulated during Dy^W^ pathogenesis are similarly perturbed in patients, including *OPN, PN, FN1, PDGFRB, CTGF, VCAN, MMP2, MMP9, TIMP1, ITGB8*, *ACTA2, CDH11, SNAI2,* and *TCF4* (Table 1), thus suggesting their relevance in pathological processes to the human condition too.

**Table 1.** Fibrosis and myofibrogenesis-related genes are upregulated in LAMA2-RD patient biopsies. Most of the genes found to be differentially expressed in LAMA2-RD patients against controls reflect the changes in expression observed in Dy^W^ mice, with the exception of *ITGB6, ILK* and *SNAI3*. Expression fold changes of up- and downregulated genes are indicated in the Log2FC column. * indicates non-significant genes with a q-value above the threshold limit of 0.05

### Treatment with the angiotensin II type 1 receptor blocker Losartan attenuates myofibrogenesis gene dysregulation

We lastly examined if the increased expression of above-mentioned growth factors, matricellular proteins, and myofibroblast markers/transcription factors in adult (7-weeks old) Dy^W^ mice could be reversed in response to an anti-fibrotic therapy, here represented by Losartan. In short, similar to data obtained for Tgf-β activating integrins and matricellular proteins, we found resolution of the dysregulation of fibrotic growth factors as well as myofibroblast drivers and markers (Figs. 8-10). These results suggest the importance of these pathways for driving fibrosis and their potential applicability as biomarkers of both fibrotic pathology and fibrotic resolution.

**Fig. 8.**
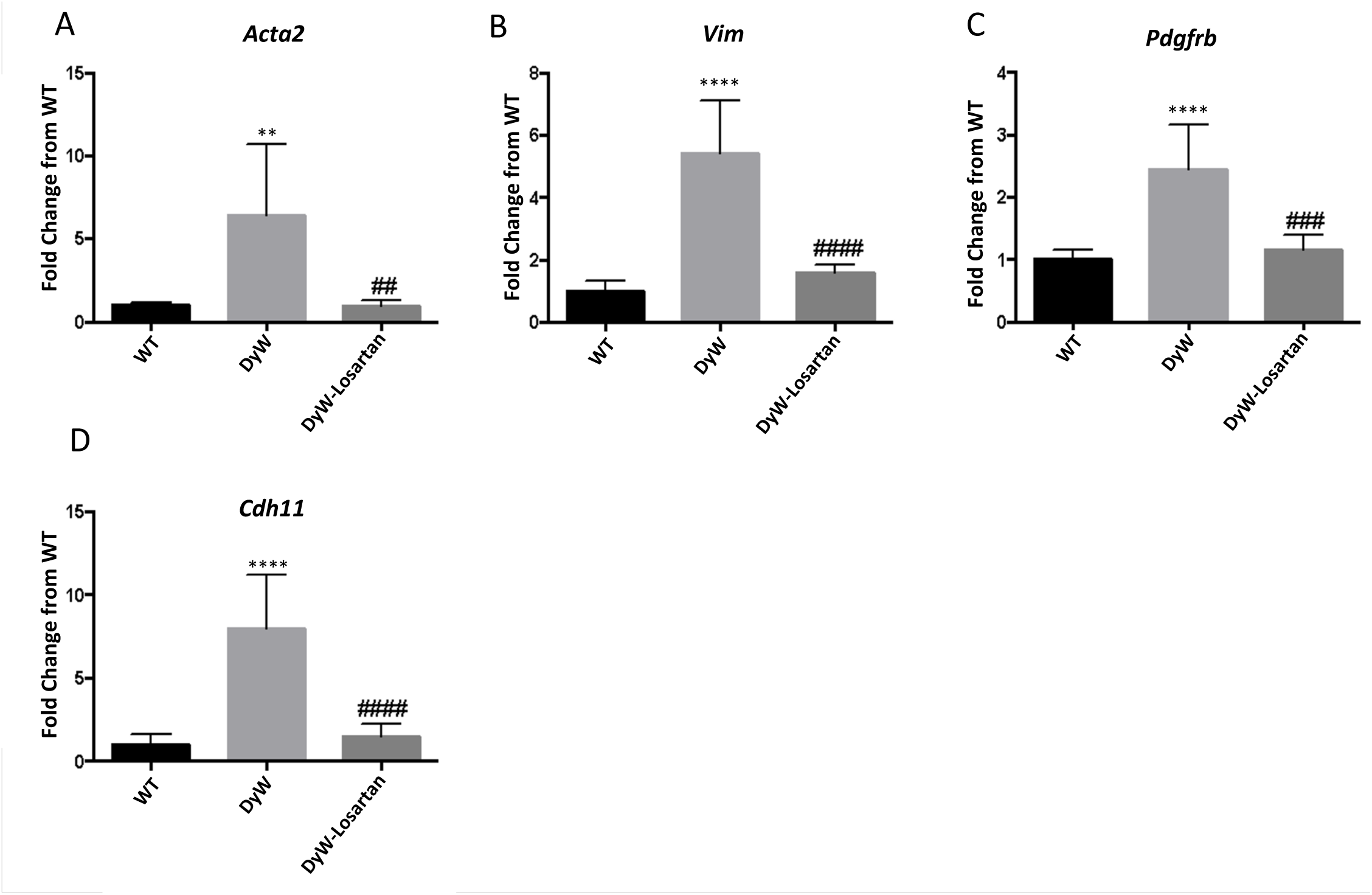
Losartan attenuates myofibroblast protein dysregulation. A-D) Gene expression measured by qRT-PCR shows significant downregulation (p<0.01, n=5, one-way ANOVA) in myofibroblast-related proteins including alpha-smooth muscle actin (A), vimentin (B), platelet derived growth factor receptor beta C), and osteoblast cadherin (D) in response to Losartan. * denotes significance from WT; # denotes significance between Dy^W^ and Dy^W^-Losartan. *=p<0.05, **=p<0.01, ***=p<0.001, ****=p<0.0001. Trend also applies to #.

**Fig. 9.**
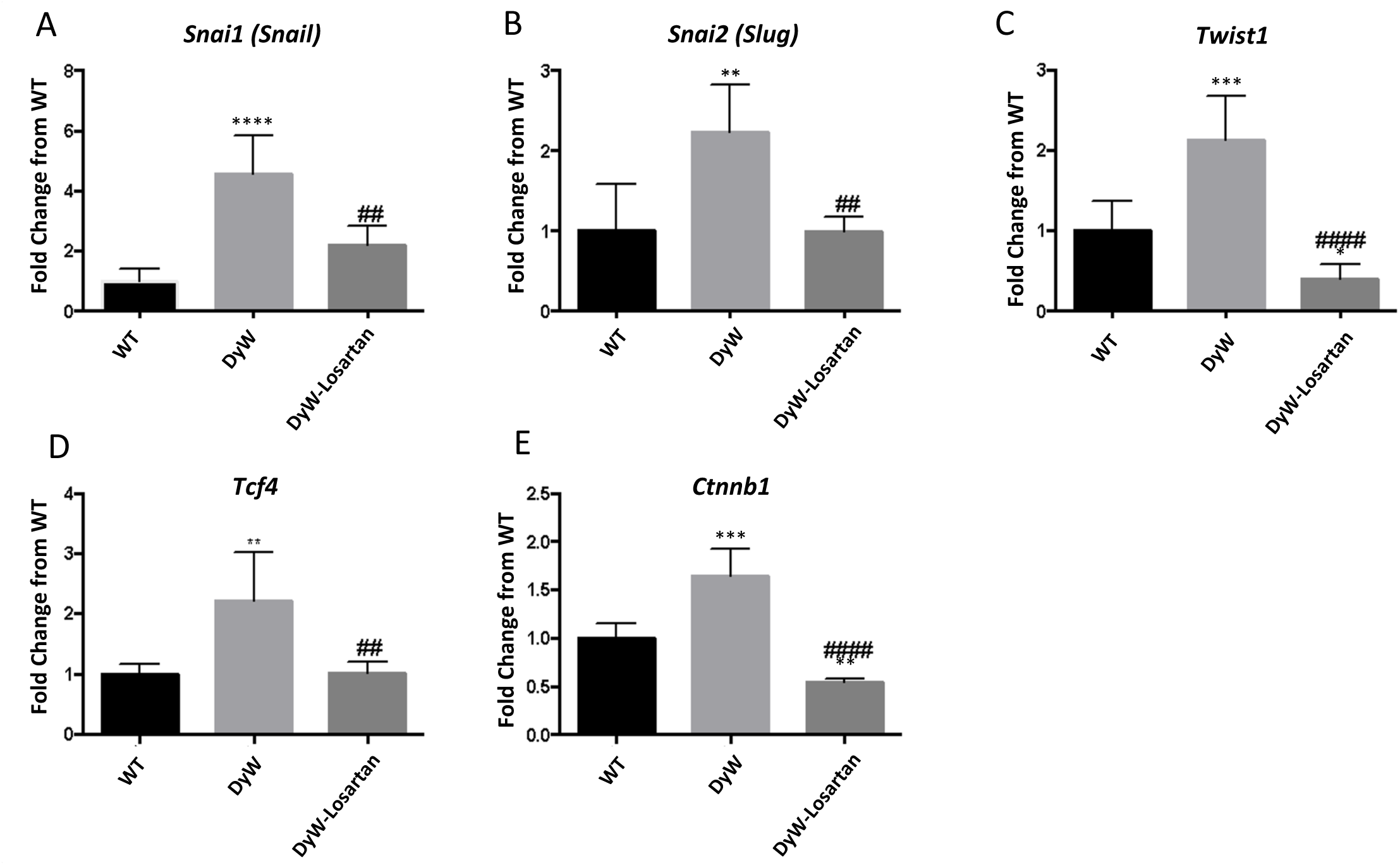
Losartan attenuates myofibroblast transcription factor dysregulation. A-E) Gene expression measured by qRT-PCR shows significant downregulation (p<0.01, n=5, one-way ANOVA) of transcription factors known to drive myofibroblast transdifferentiation including snail (A), slug (B), twist1 (C), TCF4, and β-catenin in response to Losartan. Interestingly, β-catenin and twist gene expression is significantly less in Losartan-treated mice than WT mice. * denotes significance from WT; # denotes significance between Dy^W^ and Dy^W^-Losartan. *=p<0.05, **=p<0.01, ***=p<0.001, ****=p<0.0001. Trend also applies to #.

**Fig. 10.**
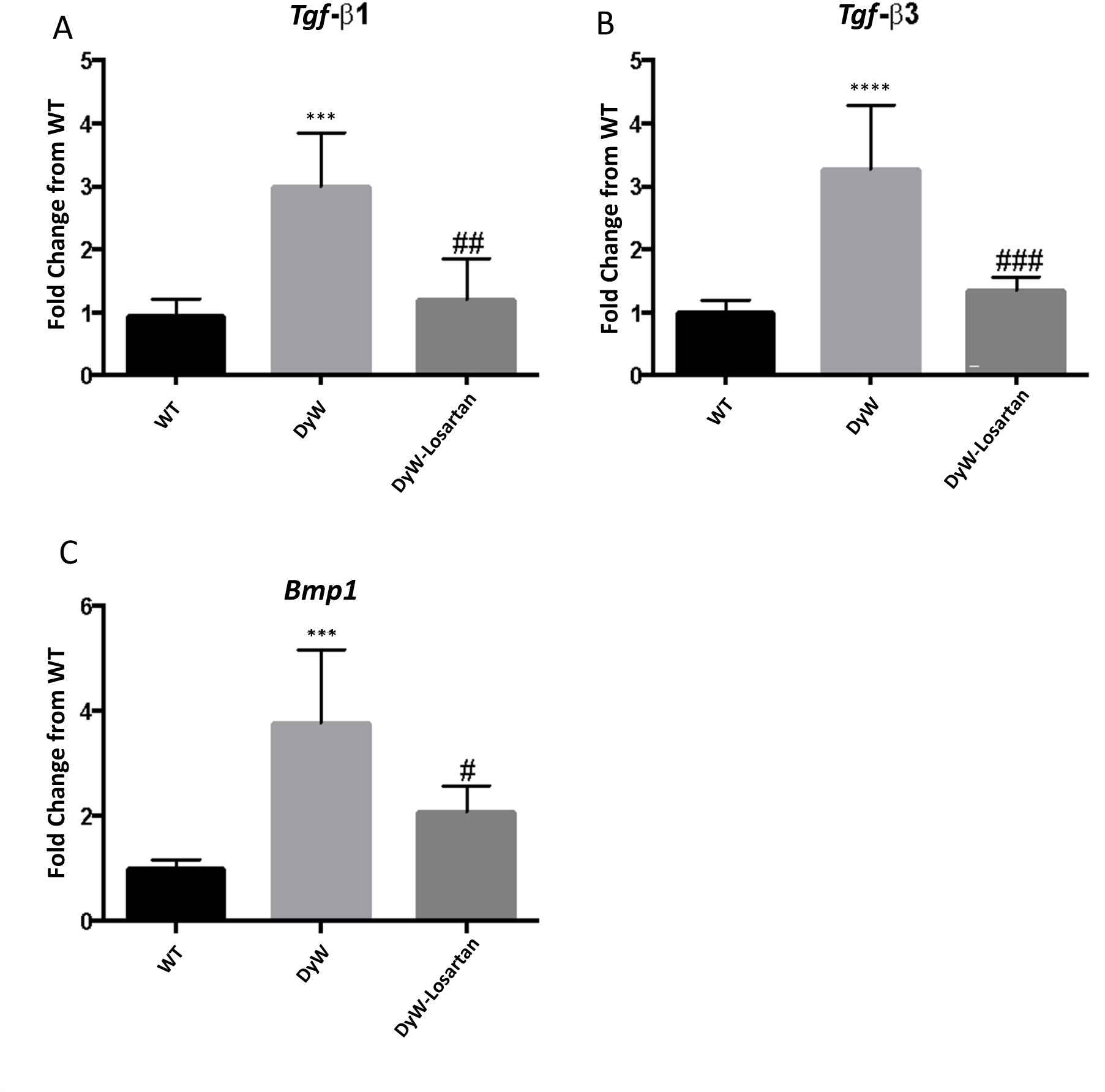
Losartan attenuates growth factor upregulation. A-C) Gene expression measured by qRT-PCR shows significant downregulation (p<0.05, n=5, one-way ANOVA) of growth factors TGF-β1 and –β3 as well as *BMP1* in response to Losartan. * denotes significance from WT; # denotes significance between Dy^W^ and Dy^W^-Losartan. *=p<0.05, **=p<0.01, ***=p<0.001, ****=p<0.0001. Trend also applies to #.

## Discussion

It is well established that integrin-αV activates TGF-β by cleaving it from the latent complex to promote fibrosis and regulate trans-differentiation of various cell types to myofibroblasts^30,33,48^. During normal wound healing, multiple cell types including resident fibroblasts as well as infiltrating cells can become activated and morph into myofibroblasts. These are ECM secreting, highly proliferative contractile cells critical to normal wound repair but can become pathological in scenarios of chronic damage and prolonged TGF-β signaling^37,42,49^. We showed here the dysregulation of integrins and their matricellular ligand partners, Tgf-β pathway and myofibrogenesis-related proteins began as early as postnatal week one in Dy^W^ mice and remain significantly elevated into adulthood. We also showed that the same pathways are also significantly elevated in human patients during a similar developmental period. Our data further demonstrates the early genesis of fibrosis, and provides a plausible explanation for the early onset muscle pathology in this disease by tipping the balance towards fibrotic matrix deposition that prevents postnatal myogenesis.

Cells from various lineages become myofibroblasts in response to signaling from the surrounding ECM milieu^31,38,40^. Integrins are instrumental for instructing cells towards adaptations promoted by matrix interactions^18,31,49,50^. Transformation of the ECM towards a pro-fibrotic micro-environment is driven by fibronectin and other matricellular proteins like osteopontin and periostin, overexpressed in Dy^W^ muscle tissue. Cues from remodelled ECM switches integrins from their low affinity “off” states to their high affinity “on” states and further induce signaling from the integrins in either outside in, i.e., traditional ligand-mediated signaling, or inside out, where adaptor proteins intracellularly dictate integrin behaviour in the ECM^17,19,22,25,26,30,51^. Extracellular signals can induce integrin-αV to bind and release TGF-β from the latent complex, initiate intracellular TGF-β signaling as well as promote Akt signaling, both of which are hyperactive in Dy^W^ muscle tissue^52^. The crosstalk of these pathways induces transcription factors like Snail, Slug, and Twist (that drive myofibroblast transition) and β-catenin, further facilitating ECM remodeling. This myomatrix transition also negatively impacts post-natal myogenesis. Together then these pathways create a circle of reciprocal regulation that can act as a double-edged sword toward exacerbating impaired muscle development and regeneration^16,17,19,22,24,30,53^.

Transdifferentiation has also been demonstrated in myoblasts. Indeed, it has been shown that C2C12 myoblasts have the ability to undergo transdifferentiation when exposed to TGF-β both *in vitro* and *in vivo*^29,32^. This suggests that myogenic precursor cells activated during post-natal myogenesis in the Dy^W^ mice may be influenced by the fibrotic microenvironment to transdifferentiate to myofibroblasts. In support of this hypothesis, the injection of recombinant TGF-β into injured skeletal muscle has been shown to induce the transdifferentiation of myoblasts to myofibroblasts^12^. Pessina *et. al* showed that, in response to chronically dysregulated TGF-β signaling in dystrophin-deficient *mdx* mice, cells of myogenic origins lose their muscle identity and gain a plasticity toward a mesenchymal-fibrogenic cell fate, thereby precluding them from participating in normal muscle regeneration^54^.

Furthermore, myofibroblast-related transcription factors (Snail, Slug, and Twist) also play a direct role in obstructing myogenesis, particularly differentiation. These transcriptions factors are part of the intricate enhancer switching process from proliferation to differentiation during myogenesis. An increased presence of these proteins prevents enhancer switching during differentiation and blocks MyoD from transcribing myogenic differentiation genes, thus inhibiting proper myogenesis^55,56^. Taken together, it is possible that the impaired post-natal growth could be due to the transitioning of satellite cells to acquire a myofibroblast-like phenotype.

We were able to demonstrate that our observations in mouse models of LAMA2-RD pathology are also relevant for human pathology as we found significant overlap in the upregulation of ECM, matricellular, and myofibroblast transdifferentiation-related genes in LAMA2-RD biopsies, confirming that not only fibrosis is a very early signature of this disease but also the EMT phenomenon can be playing a negative role in early development in these children. This could play a role in the lack of muscle mass in the children with LAMA2-RD and extensive fibrosis during early development. We did not, however, see significant upregulation integrin-αV and cognate beta partners in patient biopsies. There are a few potential reasons for this disconnect. Firstly, the control biopsies were obtained from children with presumed muscle disease but without confirmed diagnosis of muscular dystrophies which could dilute any upregulation in pathological integrins over control. Secondly, as integrin-αV is known to be a developmentally upregulated protein57, it could be more highly expressed in human development at these time-points. Lastly, and importantly as biopsies are small snapshots of human muscle, they are biased by selection. It should be noted that immuno-histochemistry in the mouse tissues showed that expression of integrin-αV was localized to only certain parts of the muscle tissue.

Finally, to tie our previous findings of Losartan-mediated attenuation of integrins and matricellular protein expression^28^, we noted that Losartan has also been shown to inhibit the expression of myofibroblast markers and transcription factors. Our results are aligned with the findings that Losartan blocked increases in α-SMA, Col1a1 mRNA, and-transdifferentiation of human lung treated with Ang II^58^. Losartan treated cardiac endothelial cells from mitral valve also resulted in down-regulation of α-SMA, Snail and Slug expression and therefore likely preventing endothelial to myofibroblast transdifferentiation (EndMT) ^59^. Anti-fibrotic therapies indeed hold significant promise for diseases such as LAMA2-RD where matrix remodeling appears to be etiological to disease progression. In addition, antifibrotic intervention may also be important for adjuvant therapy to improve efficacy of gene therapy. We and others have shown the importance of a proper myomatrix for myogenesis and it is logical to conclude that the efficiency of gene replacement may not be fully restorative in a dysregulated microenvironment. We have demonstrated that a combinatorial therapy of Losartan and *IGF-1* overexpression/ postnatal growth hormone treatment significantly enhanced muscle growth than either alone^60^. Additionally, it has been shown in *mdx* mice that combinatorial therapy of anti-fibrotic approaches with micro-dystrophin gene therapies yielded enhanced rescue compared to gene therapy alone^61^. Together, these data support the continued research and development into the etiology of matrix remodeling in muscular dystrophies and the consideration of combinatorial therapies for these complex, severe diseases.

To summarize, we confirmed here the previous results that point to integrin-αV and myofibroblast differentiation playing a likely etiological role in driving Dy^W^ pathology. Further, dysregulation of many of the proteins in human tissues lends more credence to the concept that these pathways are critical in driving pathology downstream and likely are key contributors to the insufficient postnatal myogenesis that leads to failure to thrive in LAMA2-RD patients. Lastly, based on the ability of non-specific anti-fibrotic therapies to reverse these signaling dysregulations, we believe the targets outlined in this manuscript identify key molecular players in muscle fibrosis that may be driving the disease progression and warrant further investigation for anti-fibrotic therapies in LAMA2-RD.

## Materials and methods

### Animal breeding and care

All animals were housed at the Laboratory Animal Care Facility – Charles River Campus (LACF-CRC) of Boston University on a 12:12-hr light-dark cycle. Food and water were provided *ad libitum*. All procedures were performed in accordance to the protocol approved by the IACUC of Boston University (protocol number 13-055). Heterozygous B6.129 *Lama2^dy-W/+^* mice, carrying a targeted mutation in the *Lama2* gene, were kindly provided by Dr. Eva Engvall (Burnham Institute, La Jolla, CA, USA). Genotyping was done by PCR assays on tail snips using the REDExtract-N-Amp^TM^ kit (Sigma Aldrich, St. Louis, MO) and mice homozygous for the mutation were used as the study animals (annotated Dy^W^ throughout). Losartan treatment was started at two weeks post-natal and continued until collection at seven weeks of age. Losartan was provided in the drinking water *ad libitum* (600mg/L, Cozaar by Merck pharmaceuticals) and supplemented with 25g/L of sucrose to increase palatability.

### Muscle tissue collection

Mice were first briefly bled from the submandibular vein to obtain blood samples and left to clot for 15 minutes at room temperature before being spun at 2,000 g for 8 minutes to separate serum. Animals were then euthanized with isofluorane (Webster Veterinary, Devens, MA, USA) before isolating the tibialis anterior (TA), gastrocnemius/soleus complex (GS), and quadriceps muscles (QD). Tissues were weighed and snap frozen in liquid nitrogen for RNA and protein extraction. TA/GS muscles used for histology were embedded in Tissue-Tek OCT Compound (Sakura Finetek USA, Inc., Torrance, CA, USA) and frozen in isopentane (Sigma-Aldrich, St. Louis, MO, USA) chilled in liquid nitrogen. Serial transverse sections (7μm) were prepared using the Leica CM 1850 cryostat (Leica Microsystems, Inc.) and stored at -80 °C^62^.

### Isolation and culture of muscle fibroblasts

Fibroblasts were isolated from pooled hindlimb muscles of 7-weeks old Dy^W^ and WT mice. Extracted muscles were minced in sterile HBSS in a petri dish to increase surface area exposed to pronase. Connective tissue was digested from homogenates by exposing to pronase (Sigma-Aldrich; CAT #10165921001) for 45 minutes at 37 ^0^C. The homogenate agitated vigorously by pipetting every 15 minutes to disrupt basement membrane and ECM. Following pronase incubation, muscle homogenates were filtered through 70μm and 40μm sterile filters and spun for 10 minutes at 1000 RPM. Following spin, pellet was resuspended in DMEM media supplemented with 10% FBS and 1% Pen-Strep and plated for 30 minutes on a non-coated tissue culture plate to allow fibroblasts to adhere. Media was removed (along with unadhered cells) and changed with fresh medium. Fibroblasts were cultured to 80% confluence before splitting and subsequent seeding for experimentation.

### Gene expression

RNA from ∼25 mg liquid nitrogen snap frozen pooled hind limb muscles was extracted with TRIzol reagent (Invitrogen, Carlsbad, CA) according to the manufacturer’s instructions. 1μg RNA was reverse-transcribed with the High Capacity cDNA Reverse Transcription Kit (Applied Biosystem, Foster City, CA, USA). Analysis of gene expression was performed with TaqMan assays (Applied Biosystems, Foster City, CA, USA) on ABI 7300 Real Time PCR system. 18s ribosomal subunit RNA served as the endogenous control and gene expression was calculated by using the DDCt method.

### Immunohistochemistry

Frozen tissue sections were incubated in Acetone for 20 minutes then left to air dry for 15 minutes. Protocol for Mouse on Mouse (M.O.M.) serial immunostaining for frozen sections provided by Vector Labs (Burlingame, CA) was followed using anti-Desmin antibodies (Sigma Aldrich, St. Louis, MO), anti-snail (Cell Signaling Technology, Danvers, MA, USA), anti-slug (Cell Signaling Technology, Danvers, MA, USA), anti-vimentin (Cell Signaling Technology, Danvers, MA, USA), anti-α-SMA (Sigma Aldrich, St. Louis, MO). Sections were then stained with either Alexa fluor goat anti mouse or goat anti rabbit secondary antibodies for an hour protected from light. Finally, sections were mounted with Vectashield with DAPI (Vector Labs, Burlingame, CA) or 2:1 Glycerol:PBS and imaged with a Nikon DSFi1 camera head attached to a Nikon ECLIPSE 50i light microscope system. These images were analyzed using NIS-Elements Basic Research 3.0 software.

### Patient biopsy RNA-sequencing methodology and analyses

Quadriceps muscle biopsy samples were obtained from the MRC Centre for Neuromuscular Diseases Biobank (London) or neuromuscular centers in Italy. RNA was extracted from 4 LAMA2-RD patient samples and 4 age-matched (0-3 years old) non-LAMA2-RD controls by using the RNAeasy kit (QIAgen, Ltd., Manchester, UK). RNA-sequencing was performed on 150 ng RNA at the Centre of Mendelian Genomics at the Broad Institute of MIT and Harvard, with a depth of coverage of 75 million paired-end reads. Sequencing outputs were aligned to a reference genome (hg38 Ensembl) through the STAR algorithm^63^. Differential expression analysis was run in the R programming language using the DESeq2 package^64^, that automatically performed raw reads data normalization and statistical analysis. DESeq2 outputs included a list of differentially expressed genes, the fold changes of expression indicated as base2 logarithm (or Log2FoldChange) and scores indicating the statistical significance, i.e. p-values and p-values adjusted for multiple comparisons (or q-values). Genes with a q-value equal to (or minor than) 0.05 were considered for further analysis, in accordance with DESeq2 guidelines.

### Statistics

One- and two-way ANOVA as well as student t-tests were performed using GraphPad Prism 6 software. Data are presented as mean ± standard deviation.

## Acknowledgements

Francesco Muntoni – FM is supported by the NIHR Great Ormond Street Hospital Biomedical Research Centre. The support of the NIHR, the MRC and the MDUK for the biobank at the Dubowitz Neuromuscular Centre in London is gratefully acknowledged. The views expressed in this paper are his and not necessarily those of the NHS, NIHR or the department of health.

## Author Contributions

AA completed gene expression, histological experiments, drafted/revised manuscript; AK completed cell culture experiments and performed animal husbandry; VP performed human molecular profiling analyses/contributed to manuscript revisions and editing; FM conceived human molecular profiling studies, contributed to manuscript revisions; MG conceived of overall experimental design, revised manuscript.

