## Supplementary material for "Matrix remodeling plays an etiological role in driving laminin-α2 deficient pathology": Table 1

| GENE_ID | GENE NAME | Log2FC | P_VALUE | Q_VALUE |
|---|---|---|---|---|
| COL1A1 | collagen type I alpha 1 chain | 4.11509082 | 1.84E-21 | 4.12E-18 |
| COL1A2 | collagen type I alpha 2 chain | 3.61513131 | 1.55E-24 | 6.98E-21 |
| SPP1 | osteopontin | 4.71773019 | 8.51E-07 | 2.88E-05 |
| POSTN | periostin | 3.55738058 | 1.36E-15 | 8.05E-13 |
| FN1 | fibronectin 1 | 1.82898826 | 0.00012177 | 0.00151366 |
| MMP2 | matrix metallopeptidase 2 | 1.71481398 | 6.05E-06 | 0.00013894 |
| MMP9 | matrix metallopeptidase 9 | 4.18832168 | 0.00394135 | 0.02155263 |
| TIMP1 | TIMP metallopeptidase inhibitor 1 | 1.85778075 | 1.6185E-05 | 0.00030347 |
| COL6A1 | collagen type VI alpha 1 chain | 1.32462672 | 1.5992E-06 | 4.8331E-05 |
| COL6A2 | collagen type VI alpha 2 chain | 1.4181406 | 2.75E-05 | 0.00046133 |
| COL6A3 | collagen type VI alpha 3 chain | 1.93265862 | 5.64E-06 | 0.00013083 |
| COL6A6 | collagen type VI alpha 6 chain | 2.06096827 | 4.91E-06 | 0.0001173 |
| VCAN | versican | 2.52968735 | 9.41E-08 | 4.79E-06 |
| VIM | vimentin | 1.54901054 | 1.91E-06 | 5.53E-05 |
| MCP-1 | monocyte chemoattractant protein 1 | 3.62480328 | 6.95E-10 | 8.03E-08 |
| BMP1 | bone morphogenetic protein 1 | 1.28463499 | 0.01079404 | 0.0457314 |
| CTGF | connective tissue growth factor | 3.35977213 | 1.1441E-11 | 2.3115E-09 |
| TGFB1* | transforming growth factor beta 1 | 1.05803813 | 0.05217053 | 0.15163026 |
| TGFB2 | transforming growth factor beta 2 | 0.80892179 | 0.00319056 | 0.01835982 |
| TGFB3 | transforming growth factor beta 3 | 1.70580413 | 5.1967E-06 | 0.0001228 |
| ITGA6* | integrin subunit alpha 6 | 0.48279769 | 0.22334982 | 0.41409248 |
| ITGB1* | integrin subunit beta 1 | 0.12243439 | 0.61807749 | 0.7770597 |
| ITGB3 | integrin subunit beta 3 | 1.64278115 | 0.0005419 | 0.00470649 |
| ITGB5* | integrin subunit beta 5 | 0.11177372 | 0.72253441 | 0.84804095 |
| ITGB6 | integrin subunit beta 6 | -1.42076316 | 0.00062549 | 0.00524357 |
| ITGB8 | integrin subunit beta 8 | 2.81753263 | 0.00264974 | 0.01590054 |
| ITGA5 | integrin subunit alpha 5 | 0.83280937 | 0.00294504 | 0.01728861 |
| ITGAV * | integrin subunit alpha V | 0.50440391 | 0.22981397 | 0.42199449 |
| ILK * | integrin linked kinase | -0.39400707 | 0.34227931 | 0.54815109 |
| ACTA2 | actin alpha 2, smooth muscle | 1.90786572 | 0.00703036 | 0.03316279 |
| CDHOB | osteoblast cadherin | 2.50205219 | 8.9662E-15 | 4.676E-12 |
| PDGFRB | platelet-derived growth factor receptor beta | 1.1238584 | 0.00012397 | 0.00153252 |
| SNAI2 | slug | 1.53584362 | 1.772E-05 | 0.00034254 |
| SNAI3 | snail | -1.60989122 | 1.0879E-05 | 0.00023166 |
| TWIST1 | twist | 2.49864747 | 0.00012625 | 0.00155306 |
| TCF4 | T-cell factor | 1.53755594 | 1.06E-06 | 3.46E-05 |

*Table 1-Fibrosis and myofibrogenesis-related genes are upregulated in MDC1A patient biopsies.* Most of the genes found to be differentially expressed in LAMA2-CMD patients against controls reflect the changes in expression observed in DyW mice, with the exception of ITGB6, ILK and SNAI3. Expression fold changes of up- and downregulated genes are indicated in the Log2FC column. * indicates non-significant genes with a q-value above the threshold limit of 0.05.
